# Refolding of lid subdomain of SARS-CoV-2 nsp14 upon nsp10 interaction releases exonuclease activity

**DOI:** 10.1101/2022.02.17.480845

**Authors:** Anna Czarna, Jacek Plewka, Leanid Kresik, Alex Matsuda, Abdulkarim Karim, Colin Robinson, Sean O’Byrne, Fraser Cunningham, Irene Georgiou, Magdalena Pachota, Grzegorz Popowicz, Paul Graham Wyatt, Grzegorz Dubin, Krzysztof Pyrć

**Affiliations:** Virogenetics Laboratory of Virology, Malopolska Centre of Biotechnology, Jagiellonian University, Gronostajowa 7a, 30-387 Krakow, Poland; Department of Biology, College of Science, Salahaddin University-Erbil, Kirkuk Road, 44002 Erbil, Kurdistan Region, Iraq and Department of Community Health, College of Health Technology, Cihan University-Erbil, 100 street, 44001 Erbil, Kurdistan Region, Iraq; Drug Discovery Unit, Wellcome Centre for Anti-Infectives Research, School of Life Sciences, University of Dundee, Dow Street, Dundee DDI 5EH, U.K.; Protein Crystallography Research Group, Malopolska Centre of Biotechnology, Jagiellonian University, Gronostajowa 7a, 30-387 Krakow, Poland; Helmholtz Zentrum München, Ingolstädter Landstrasse 1, 85764 Neuherberg, Germany; Bavarian NMR Center, Department of Chemistry, Technical University of Munich, Lichtenbergstrasse 4, 85748 Garching, Germany

## Abstract

During the RNA replication, coronaviruses require proofreading to maintain the integrity of their large genomes. Nsp14 associates with viral polymerase complex to excise the mismatched nucleotides. Aside from the exonuclease activity, nsp14 methyltransferase domain mediates cap methylation, facilitating translation initiation and protecting viral RNA from recognition by the innate immune sensors. The nsp14 exonuclease activity is modulated by a protein co-factor nsp10. While the nsp10/nsp14 complex structure is available, the mechanistic basis for nsp10 mediated modulation remains unclear in the absence of nsp14 structure. Here we provide a crystal structure of nsp14 in an apo-form. Comparative analysis of the apo- and nsp10 bound structures explain the modulatory role of the co-factor protein. Further, the structure presented in this study rationalizes the recently proposed idea of nsp14/nsp10/nsp16 ternary complex.

## Introduction

Upon release into the cytoplasm of the host cell, the coronavirus genome is translated into a single non-functional polyprotein 1a/1ab. Its autoproteolytic processing releases a number of nonstructural proteins (nsp) responsible for viral replication. The activity of particular nsp’s is relatively well-characterized, but the mechanistic understanding of their interplay is still insufficient^1–3^.

Nsp14 consists of two domains, each with a distinct catalytic activity. The C-terminal domain is a SAM-dependent methyltransferase, while the N-terminal ExoN domain exhibits 3’-5’ exonuclease activity. Recognition of unmodified RNA molecules constitutes a universal antiviral strategy embedded in the host innate immune defences. To avoid detection, coronaviral RNA is capped by viral enzymes, what additionally increases its stability and facilitates translation initiation. Capping involves a series of enzymatic reactions. In the penultimate step, the C-terminal domain of nsp14 catalyzes N7-methylation of a GpppA intermediate. Subsequent methylation by nsp16 results in a functional cap (^7Me^GpppA_2’OMe_)^4–9^.

The large coronaviral genome requires replication fidelity to maintain its functionality. While the nsp12-centred polymerase complex is highly processive, it is also error-prone. An unaudited replication would result in excessive accumulation of mutations and the generation of defective progeny virions. Nsp14 associates with the replication complex and removes incorrectly incorporated nucleotides from the 3’ end of the newly formed RNA strand. Nsp14 is associated with nsp10, a protein co-factor that modulates nsp14 exonuclease activity in the replication complex^6, 10^.

The structural basis of nsp14 interaction with nsp10 has been elucidated^11^. We have recently demonstrated that nsp14, nsp10 and nsp16 form a ternary complex, further modulating nsp14 catalytic activity^12^. Despite the detailed structural characterization of the complexes, the mechanistic basis of the modulation of nsp14 exonuclease activity remains elusive. The absence of structural information on nsp14 alone hinders the mechanistic understanding of the consequences of binding with nsp10 and nsp10/nsp16.

Here, we provide the crystal structure of nsp14 in the apo-form (without nsp10), which, together with prior data on nsp10/nsp14 complex, provide insight into the modulatory role of nsp10 on nsp14 exonuclease activity. Furthermore, the presented structure strongly supports the nsp14/nsp10/nsp16 heterotrimer formation model.

## Results and discussion

Full length SARS-CoV-2 nsp14 exonuclease/methyltransferase was expressed in *E. coli* and purified. The crystal structure of S-adenosylhomocysteine (SAH) bound nsp14 was determined at 2.5 Å resolution (**Table I**). The crystals belonged to P21 space group and contained 2 molecules in the asymmetric unit. Nsp14 is characterized by a bimodular structure corresponding to the two distinct catalytic activities (**Figure 1A**). The C-terminal methyltransferase domain of nsp14 (amino acids 287-524) is well defined by the electron density for most of its part safe for amino acids 454-465. At the same time, a significant portion of the N-terminal exonuclease domain (amino acids 25-286) is undefined by the electron density, indicating significant flexibility. This, in particular, relates to amino acids 40-44, 93-103, and 123-154, and is of importance for further discussion. The undefined parts are identical in both chains.

**Figure 1.**
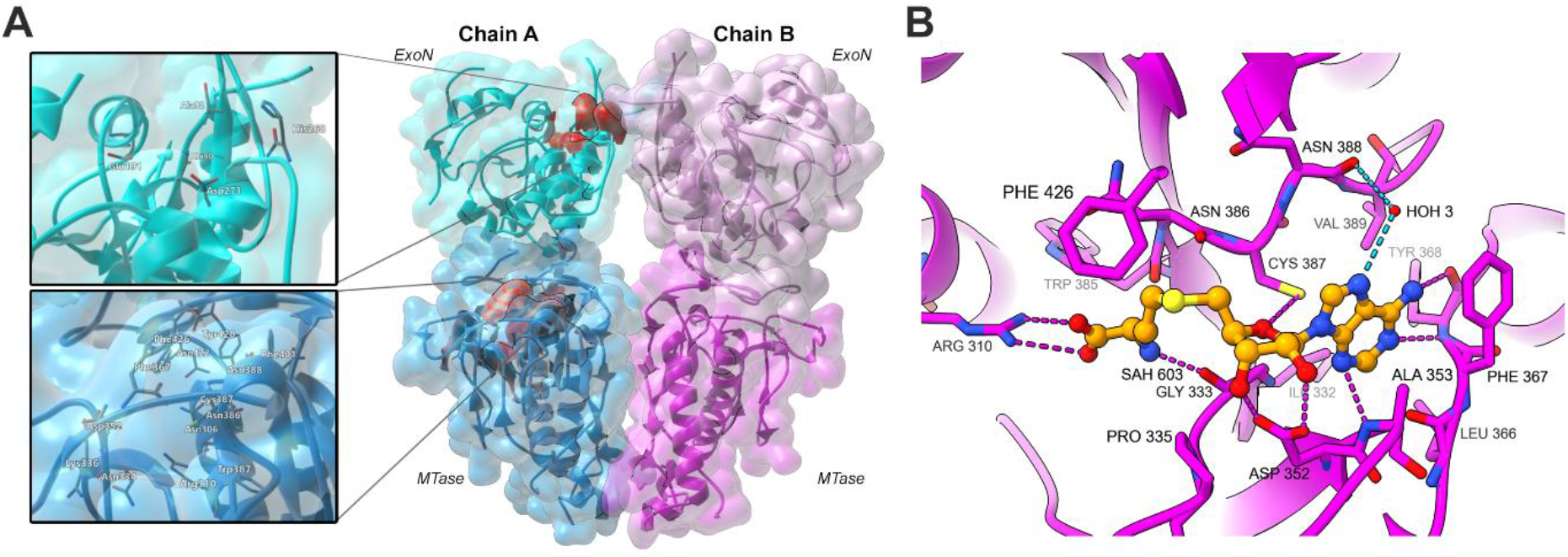
Crystal structure of apo-nsp14. (A) Arrangement of the molecules in the asymmetric unit. Model A – blue, model B – pink. MTase and ExoN domains within the bimodular structure of nsp14 are distinguished by primary colour shade. Active site residues are indicated in red in one molecule, and the detailed arrangements of the active sites are shown in inserts. (B) Interactions of SAH (orange, stick model) at the active site of nsp14 MTase domain (pink, ribbon). Hydrogen bonds are depicted as dotted lines; waters are indicated as red spheres.

**Table I.** Data collection and refinement statistics. Values provided in parentheses are for the highest resolution shell.

| Data collection statistics |  |
| --- | --- |
| PDB ID | 7R2V |
| Wavelength (Å) | 1.00004 |
| Resolution range | 86.59 - 2.53 |
| Space group | P 1 21 1 |
| Unit cell | 67.18, 100.31, 90.87 (Å) |
|  | 90.00, 107.66, 90.00 (°) |
| Data scaling software | autoPROC (Version 1.0.5),<br>AIMLESS, STARANISO |
| Total reflections | 19,046 |
| Unique reflections | 19,046 |
| Multiplicity | 4.62 (4.72) |
| Completeness (%) | 100.00 (99.92) |
| Mean I / sigma (I) | 3.80 (0.33) |
| Wilson B-factor (Å<sup>2</sup>) | 41.31 |
| R-merge | 0.2117 (1.5567) |
| CC1/2 | 0.9510 (0.1646) |
| CC* | 1 (0.962) |
| Structure refinement statistics |  |
| Total reflections | 19,046 |
| Unique reflections | 19,046 |
| Wilson B-factor | 41.31 |
| Reflections in refinement | 19,045 |
| Reflections in free set | 911 |
| R<sub>work</sub> | 0.201 |
| R<sub>free</sub> | 0.265 |
| FSC average | 0.8907 |
| RMSD bonds | 0.0056 |
| RMSD angles | 1.418 |
| Ramachandran favoured (%) | 90.4 |
| Ramachandran allowed (%) | 8.8 |
| Ramachandran outliers (%) | 0.8 |
| Rotamer outliers | 7.6 |
| Clash score | 4.9 |
| Overall number of atoms (non-H) | 7,061 |
| in macromolecules | 6,993 |
| in ligands | 20 |
| in solvent | 48 |
| Average B-factor | 56.4 |
| for macromolecules | 56.7 |
| for ligands | 44.4 |
| for solvent | 29.4 |

The methyltransferase domain of nsp14 is characterized by a fold atypical to methyltransferases. The structure is built around a central β-sheet composed of 5 instead of 7 canonical β-strands. Further, the methyltransferase domain of nsp14 contains an atypical peripheral zinc finger stabilized by another element that distinguishes nsp14 from the typical methyltransferase fold, a supplementary C-terminal α-helix. The methyltransferase fold within our structure closely resembles that previously seen in the nsp10/nsp14 complex of SARS-CoV-1 (PDB ID 5C8S) with rmsd of 0.57Å for 197 equivalent C_α_ atoms. A reaction product, SAH, is clearly defined by the electron density in our structure (**Figure 1B**), but otherwise, the active site is empty (the methylated reaction product is absent). Interestingly, the active site faces a large, unobstructed water channel. As such, our structure is particularly valuable in terms of inhibitor development (fragment screening and inhibitor soaking).

The SAM/SAH-binding region in our structure (331-DxGxPxA-337) diverges from the classical methyltransferase motif I (E/DxGxGxG; ^13^) but is consistent with that described in nsp14 of SARS-CoV-1 ^14^. The purine ring of SAH is positioned *via* sulphur/π interaction with Cys387; hydrophobic interactions contributed by sidechains of Ala353, Val389, Phe367, and direct hydrogen bonds contributed by mainchain of Tyr368. A water-mediated hydrogen bond contributed by Asn388 is additionally observed in model B. Ribose is stabilized by hydrogen bonds contributed by the sidechain of Asp352. Mainchain carbonyl oxygens of Gly333 and Trp385 contribute hydrogen bonds with the cysteine amine, whereas its carboxyl group forms a salt bridge with Arg310. The identified SAH positioning residues are consistent with the conclusions of prior mutagenesis studies ^15^. The majority of reported nanomolar MTase inhibitors are SAM competitive ^16^. By delineating the detailed molecular interactions in the SAH pocket, our study facilitates the development of SAH mimetic inhibitors of SARS-CoV-2 MTase.

A structure of nsp14 from SARS-CoV-2 has been recently deposited at the protein data bank (PDB ID: 7QGI), which contains an empty active site (no substrates or reaction products). Comparison with our structure demonstrates that the SAM/SAH binding site in 7QGI is occluded by a loop connecting β2’ and β3’ strands of a central β-sheet. We initially thought this could represent a mechanism of reaction product removal or a methyltransferase activity regulatory mechanism. However, a closer look at 7QGI crystal structure demonstrates that the place where the loop in question is located in SAH bound structures (nsp10/nsp14 complex from SARS-CoV-1 and this study) is occluded by an adjacent molecule in 7QGI. This suggests that the blocked active site in 7QGI is an artefact of crystal packing rather than representing a physiologically relevant conformation. Supporting the above conclusion, when Ma and colleagues^14^ analyzed SARS-CoV-1 nsp10/nsp14 complexes in SAH bound and empty-forms, the loop in question had the same conformation in both SAH bound and empty structure (and comparable to that found in our structure), further suggesting that 7QGI represents a non-physiological conformation within the discussed feature.

The exonuclease domain of nsp14 is built around a five-stranded, twisted β-sheet flanked by α-helices. Such arrangement of the core structure elements follows the overall design of DEDD family exonucleases but contains unique features: the N-terminal region (1-84) and the first zinc finger. The striking features of the exonuclease domain in our structure became evident only upon comparison with the structure of nsp10/nsp14 complex. The N-terminal of nsp14 spanning amino acids 1-70 is folded and oriented differently in compared structures (**Figure 2A, B**). The N-terminal is well defined by electron density in the nsp10/nsp14 complex, but contains little ordered secondary structures. Amino acids 1-56 constitute the first of the two interaction regions with nsp10 (nomenclature of structural features after Ma *et al.*^14^). The second interaction region comprises residues 60-67 and 200-202. In apo-nsp14 structure, residues 1-24 are not defined by the electron density. Region 25-70 is completely refolded in apo-nsp14 structure compared to the nsp10/nsp14 complex. For example, residues Thr50-Met58 form an α-helix in apo-nsp14 structure, whereas residues 51-54 form a β1-strand within β1, β5, β6 sheet in the nsp10/nsp14 complex. The remaining residues (25-49, 59-67) contribute completely different intermolecular interactions within apo-nsp14 structure compared to nsp10/nsp14 complex. In the absence of nsp10, the N-terminal region of nsp14 covers the second nsp10 interaction region, burying, among others, residues 200-202. Thereby, the N-terminus of nsp14 acts as a lid, covering the nsp10 binding site in the absence of the protein co-factor. The lid rearranges its structure in the presence of nsp10 to create a significant interaction surface with nsp10.

**Figure 2.**
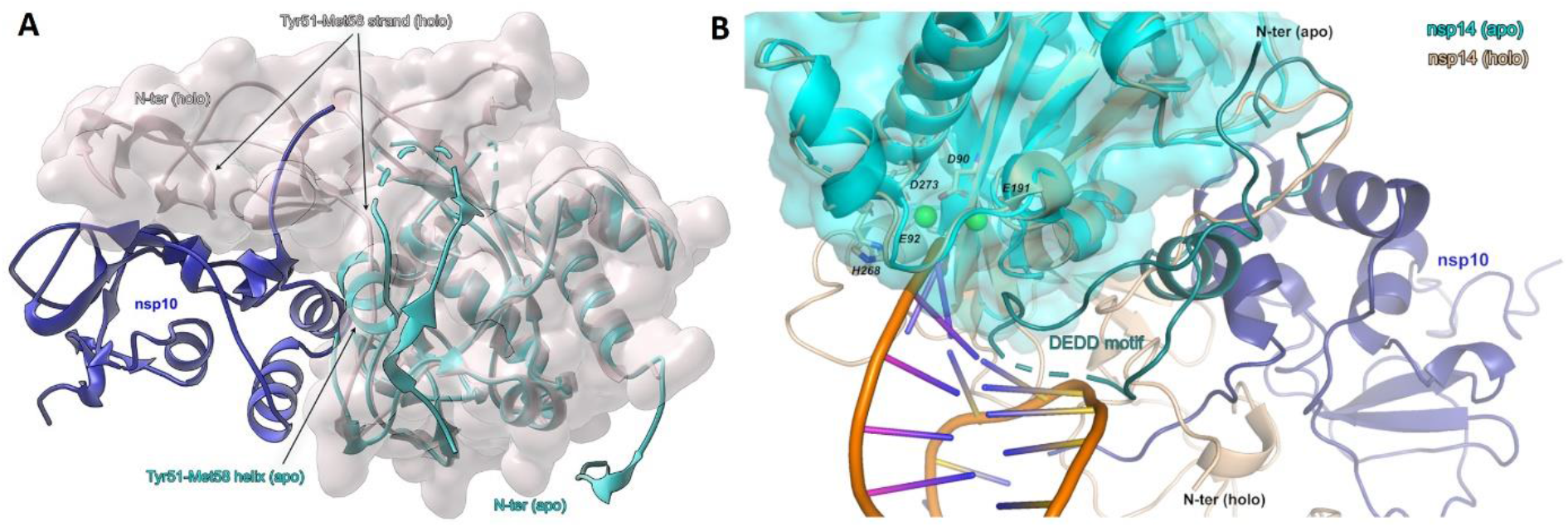
Nsp10 induced refolding of nsp14 ExoN lid subdomain. (A) Nsp10 (navy blue, ribbon) / Nsp14 (grey, surface) complex overlaid on apo-nsp14 (cyan). The N-terminal lid of nsp14 refolds upon binding of nsp10 as exemplified by Thr50-Met58 region assuming a helical structure in apo-nsp14 and forming a strand of a β-sheet in nsp10/nsp14 complex. In the apo structure, the lid occludes nsp10 binding site at the surface of nsp14. (B) The RNA interaction site of nsp10 (navy blue) /nsp14 (beige) complex (7N0B) is sterically occluded by lid subdomain in apo-nsp14 structure (cyan), providing a rationale for the negligible exonuclease activity of nsp14 in the absence of nsp10. RNA (orange). The apo-nsp14 loop, occluding the RNA binding site, is denoted in dark cyan, dotted line.

Nsp10 enhances the 3’-exonucleolytic activity of nsp14 but does not significantly affect the methyltransferase activity. Nsp10 effect on nsp14 was rationalized by nsp10/nsp14 crystal structure, where nsp10 contributes significant contacts with ExoN domain of nps14 but does not interact with the N7-methyltransferase domain^14^. The extensive surface area of interaction suggested that nsp10 maintains the structural stability of ExoN, but the nature and extent of instability of apo-nsp14 were not known in the absence of the structure. In the current study, we show that the ExoN domain of nsp14 is partially unfolded in the absence of nsp10. The proximal part of β2-β3 sheet adjacent to catalytic Glu92 is undefined by electron density (flexible, unstructured) in apo-nsp14. The same is true for the entire region between β4 and α3. Residues constituting β5-β6 and α2 in the nsp10/nsp14 complex structure are not defined by the electron density in the apo-nsp14 structure. Significantly, in apo-nsp14, the catalytic metal ion binding site within DEDD motif is partially occluded by the sidechain of His188. Further, the RNA binding site is partially occluded by the lid fragment (Thr35-Lys47), which buries the nsp10 binding site. Only nsp10 induced refolding of the lid removes the steric occlusion and induces a catalytically capable conformation of nsp14 observed in nsp10/nsp14 complex (**Figure 2B**).

We have demonstrated earlier that nsp14, nsp10 and nsp16 form a ternary complex cantered around nsp10 ^12^. However, such complex is not supported by the structural data available on nsp10/nsp14 and nsp10/nsp16 binary complexes. When the binary complexes are overlayed on nsp10, the lid of nsp14 clashes with nsp16, demonstrating that ternary complex formation would require a structural rearrangement within the lid. The current study demonstrates that the lid of nsp14 is capable of structural morphing, indirectly supporting the mechanism of triplex formation.

## Conclusions

The current pandemics highlighted the importance of understanding coronavirus physiology. Prior research on other human and animal coronaviruses facilitated an unprecedented pace of vaccine and antiviral development, but the constant genetic drift and the threat of new zoonotic transmission warrant continued effort.

Nsp14 constitutes a valid target in SARS-CoV-2, but the structural information was incomplete. This study provided the structural basis for a mechanistic explanation of nsp10 mediated modulation of nsp14 3’-exonuclease activity, defined the structure of nsp14 N7-methyltransferase in the absence of RNA substrate, and offered crystallization conditions suitable for soaking of small molecules at the active site of nsp14 N7-methyltransferase. Thereby, the study contributes to our understanding of coronavirus physiology and facilitates the effort of anti-coronaviral drug development.

## Materials and methods

### Expression construct

Sequence encoding amino acids 5926 – 6452 of SARS CoV-2 polyprotein 1ab (Uniprot P0DTD1), including D90A and D92A substitutions ^5^, flanked by BamHI and NotI cleavage sites and containing a TEV protease cleavage site was codon-optimized for expression in *Escherichia coli* and obtained by synthesis at GeneArt. The sequence was subcloned into pETDuet-1 expression vector.

### Protein expression and purification

The plasmid was transformed into *E. coli* BL21 Rosetta2 (DE3) strain. Expression was carried out in Terrific Broth medium (Bioshop) supplemented with 100 μg/ml ampicillin (Sigma-Aldrich). Bacterial cultures were incubated at 37°C until the OD_600_ value reached 1.2 - 1.4. The cultures were cooled to 4°C for 20 min, and the recombinant protein expression was induced with 0.5 mM isopropyl-D-1-thiogalactopyranoside (IPTG; Sigma-Aldrich). The bacteria were incubated at 18°C for additional 18 hours. Bacterial cells were harvested by centrifugation at 7,000 × g for 30 min at 4°C and stored at −20 frozen °C before the purification.

Bacterial pellets were resuspended in lysis buffer (50 mM Tris-HCl pH 8.5, 500 mM NaCl, 5 mM MgCl_2_, 10% v/v glycerol, 5 mM β-mercaptoethanol, 10 mM imidazole), supplemented with benzonase (Millipore) and protease inhibitors cocktail (Roche), and lysed by sonication at 80% amplitude for 20 min at 14°C with 3 s on / 2 s off cycles. The fraction of the supernatant containing soluble protein was separated from debris by centrifugation at 25,000 × g for 40 min at 4°C. The clarified supernatant was incubated overnight at 4°C with HisPur Ni-NTA Resin (Thermo Scientific) pre-equilibrated with the lysis buffer. Resin was sequentially washed with lysis buffer, then with wash buffer 1 (50 mM Tris-HCl pH 8.5, 500 mM NaCl, 5 mM MgCl_2_, 10% v/v glycerol, 5 mM β-mercaptoethanol, 20 mM imidazole) and wash buffer 2 (50 mM Tris-HCl pH 8.5, 500 mM NaCl, 5 mM MgCl_2_, 10% v/v glycerol, 5 mM β-mercaptoethanol, 30 mM imidazole). The immobilized protein was eluted with elution buffer (50 mM Tris-HCl pH 8.5, 250 mM NaCl, 5 mM MgCl_2_, 10% v/v glycerol, 5 mM β-mercaptoethanol, 250 mM imidazole). Fractions containing the protein of interest (as determined by SDS-PAGE) were pooled and supplemented with tobacco etch virus (TEV) protease in a 1:10 molar ratio. TEV cleavage was combined with overnight dialysis into a cleavage buffer (50 mM Tris-HCl pH 8.5, 200 mM NaCl, 5 mM MgCl_2_, 10% v/v glycerol, 5 mM β-mercaptoethanol, 10 mM imidazole). Reverse chromatography on HisPur Ni-NTA resin pre-equilibrated with the dialysis buffer allowed the cleaved His-tag removal. The protein was further purified by size exclusion chromatography on HiLoad 26/600 Superdex 200 prep grade column (GE Healthcare) equilibrated with the working buffer (50 mM Tris-HCl pH 8.5, 150 mM NaCl, 5 mM MgCl_2_, 1 mM TCEP). The purity of the sample was assessed by SDS-PAGE.

### Crystallization and data collection

The purified protein was concentrated to 11.5 mg/ml. The sample was supplemented with 2mM SAM. Crystallization screening was performed at 4°C using the sitting drop vapour diffusion method. Crystals were obtained from a solution containing 15 % v/v 2-propanol, 0.2 M imidazole pH 7.6 and 500 mM polypropylene glycol 400. Further optimization was performed around initial conditions where the crystals generally appeared after 4 days. The crystals were cryoprotected in 30% ethylene glycol in the mother liquor and flash-cooled in liquid nitrogen. The diffraction data were collected at Swiss Light Source (SLS, Villigen, Switzerland) beamline X06DA - PXIII. The data were indexed and integrated using XDS ^17^. The data were scaled and merged using STARANISO webserver. A molecular replacement solution was found using Phaser ^18^ and the structure of SARS-CoV-1 nsp14 (PDB code: 5C8S) as a search model. SAH was identified in the electron density map using CheckMyBlob server (https://checkmyblob.bioreproducibility.org). Restraints were obtained from ligand builder in Coot ^19^. The initial model was subjected to several iterations of manual and automated refinement cycles using COOT and REFMAC5, respectively ^20, 21^. Throughout the refinement, 5%of the reflections were used for cross-validation analysis, and the behaviour of Rfree was employed to monitor the refinement strategy.

### Data analysis

The final model was analyzed and the graphics was prepared using PyMOL (Schrödinger, LLC).

## Acknowledgements

This work was supported by a subsidy from the Polish Ministry of Science and Higher Education for research on SARS-CoV-2 and a grant from the National Science Center (UMO-2017/27/B/NZ6/02488; to KP).

## Supporting Information

### Crystal structure of apo-nsp14 in the absence of nsp10

#### Materials and Methods

##### Nsp14 construct used in the study

The initial glycine residue is a part of TEV recognition site and remains appended to nsp14 sequence in the purified nsp14.

Nucleotide sequence:

~~~
(ggc)atggctgaaaatgtaacaggactctttaaagattgtagtaaggtaatcactgggttacatcct
acacaggcacctacacacctcagtgttgacactaaattcaaaactgaaggtttatgtgttgacatacc
tggcatacctaaggacatgacctatagaagactcatctctatgatgggttttaaaatgaattatcaag
ttaatggttaccctaacatgtttatcacccgcgaagaagctataagacatgtacgtgcatggattggc
ttcgctgtagctgggtgtcatgctactagagaagctgttggtaccaatttacctttacagctaggttt
ttctacaggtgttaacctagttgctgtacctacaggttatgttgatacacctaataatacagattttt
ccagagttagtgctaaaccaccgcctggagatcaatttaaacacctcataccacttatgtacaaagga
cttccttggaatgtagtgcgtataaagattgtacaaatgttaagtgacacacttaaaaatctctctga
cagagtcgtatttgtcttatgggcacatggctttgagttgacatctatgaagtattttgtgaaaatag
gacctgagcgcacctgttgtctatgtgatagacgtgccacatgcttttccactgcttcagacacttat
gcctgttggcatcattctattggatttgattacgtctataatccgtttatgattgatgttcaacaatg
gggttttacaggtaacctacaaagcaaccatgatctgtattgtcaagtccatggtaatgcacatgtag
ctagttgtgatgcaatcatgactaggtgtctagctgtccacgagtgctttgttaagcgtgttgactgg
actattgaatatcctataattggtgatgaactgaagattaatgcggcttgtagaaaggttcaacacat
ggttgttaaagctgcattattagcagacaaattcccagttcttcacgacattggtaaccctaaagcta
ttaagtgtgtacctcaagctgatgtagaatggaagttctatgatgcacagccttgtagtgacaaagct
tataaaatagaagaattattctattcttatgccacacattctgacaaattcacagatggtgtatgcct
attttggaattgcaatgtcgatagatatcctgctaattccattgtttgtagatttgacactagagtgc
tatctaaccttaacttgcctggttgtgatggtggcagtttgtatgtaaataaacatgcattccacaca
ccagcttttgataaaagtgcttttgttaatttaaaacaattaccatttttctattactctgacagtcc
atgtgagtctcatggaaaacaagtagtgtcagatatagattatgtaccactaaagtctgctacgtgta
taacacgttgcaatttaggtggtgctgtctgtagacatcatgctaatgagtacagattgtatctcgat
gcttataacatgatgatctcagctggctttagcttgtgggtttacaaacaatttgatacttataacct
ctggaacacttttacaagacttcag
~~~

Amino acid sequence

~~~
(G)MAENVTGLFKDCSKVITGLHPTQAPTHLSVDTKFKTEGLCVDIPGIPKDMTYRRLISMMGFKMNY
QVNGYPNMFITREEAIRHVRAWIGFAVAGCHATREAVGTNLPLQLGFSTGVNLVAVPTGYVDTPNNTD
FSRVSAKPPPGDQFKHLIPLMYKGLPWNVVRIKIVQMLSDTLKNLSDRVVFVLWAHGFELTSMKYFVK
IGPERTCCLCDRRATCFSTASDTYACWHHSIGFDYVYNPFMIDVQQWGFTGNLQSNHDLYCQVHGNAH
VASCDAIMTRCLAVHECFVKRVDWTIEYPIIGDELKINAACRKVQHMVVKAALLADKFPVLHDIGNPK
AIKCVPQADVEWKFYDAQPCSDKAYKIEELFYSYATHSDKFTDGVCLFWNCNVDRYPANSIVCRFDTR
VLSNLNLPGCDGGSLYVNKHAFHTPAFDKSAFVNLKQLPFFYYSDSPCESHGKQVVSDIDYVPLKSAT
CITRCNLGGAVCRHHANEYRLYLDAYNMMISAGFSLWVYKQFDTYNLWNTFTRLQ
~~~

